# Altered Placental [1-^13^C]Pyruvate Metabolism in Spontaneous Intrauterine Growth Restricted near-term Pregnant Guinea Pigs

**DOI:** 10.64898/2026.01.23.701103

**Authors:** Lindsay E. Morris, Lanette J. Friesen-Waldner, Trevor P. Wade, Barbra de Vrijer, Timothy R.H. Regnault, Charles A. McKenzie

## Abstract

IUGR is associated with increased risk of fetal compromise, yet can be difficult to detect and phenotype with routine clinical surveillance. Given placental insufficiency and hypoxia can remodel placental energy metabolism, methods that directly assess placental metabolic function may improve identification and phenotyping of growth-restricted pregnancies. We combined structural, body composition, and hyperpolarized metabolic MRI to characterize a near-term spontaneous IUGR (spIUGR) phenotype in the guinea pig. Twenty-two pregnant guinea pig sows (71 fetuses) underwent ^1^H and hyperpolarized ^13^C MRI at 60 ± 1 days gestation to quantify fetal and placental volumes, maternal/fetal body composition, and quantify placental pyruvate metabolism. Fetuses were classified as spIUGR when ≥ 3 of 5 established markers were present (body weight, brain–body, brain–liver, brain–placenta ratios, and body weight relative to pregnancy mean); corresponding volume cut-offs were derived from weight cut-offs. MRI-derived fetal and placental volumes correlated strongly with collection weights and classified spIUGR consistently with weight-based criteria. Maternal adiposity (subcutaneous and visceral) was inversely associated with fetal adipose tissue volume, and maternal visceral fat PDFF was negatively associated with fetal adipose PDFF. Hyperpolarized MRI demonstrated IUGR phenotype-dependent placental pyruvate routing: LPR increased with asymmetric (brain-sparing) growth, showing a positive association with brain–body volume ratio (p = 0.02) and brain–body weight ratio (p = 0.04). In contrast, BPR was not significantly related to brain–body ratios (p = 0.09–0.11) but was inversely associated with absolute fetal size (body volume and weight, p = 0.02). These findings validate MRI volumetry for non-invasive identification of placental insufficiency spIUGR and link growth restriction severity and asymmetry to distinct placental metabolic signatures measurable in vivo.

## Introduction

Intrauterine growth restriction (IUGR), a condition where the fetus does not reach its full growth potential, is frequently underpinned by chronic placental hypoxia, affects about 5–10% of pregnancies and remains a key factor in perinatal morbidity and mortality, as well as adverse postnatal metabolic health [1–3]. Many IUGR cases are classified as late-onset, occurring after 32 weeks of gestation, and are often underdiagnosed because of their subtle clinical presentation [4–7]. Recent research suggests that up to 50% of late-onset IUGR cases go undetected before delivery largely because umbilical artery Doppler indices may remain within normal ranges and fetal size can appear reassuring on standard biometric assessment [5,6,8]. This pattern is consistent with uteroplacental hypoxia and adaptive placental vascular remodeling that preserves umbilical end-diastolic flow, reducing Doppler sensitivity. Accordingly, late-onset IUGR often presents with placental insufficiency and cerebral redistribution (brain-sparing), reflected by disproportionate slowing of abdominal growth relative to head growth rather than marked reductions in estimated fetal weight [3]. Despite these generally normal ultrasound results, late-onset term IUGR is linked to higher risks of fetal compromise, postnatal metabolic and neurodevelopmental impacts, and a specific neonatal morbidity profile that varies depending on the timing and severity of growth restriction and the gestational age at birth [4,9–11]. These higher risks for late-onset IUGR emphasise the need for more sensitive, functionally targeted diagnostic methods [12].

Placental hypoxia triggers a metabolic reprogramming in the placenta, in which HIF-1–mediated changes of mitochondrial oxygen utilization are thought to promote increased glycolysis and lactate generation, a Warburg-like phenotype [13–16]. The resulting shift in placental energy metabolism, characterized by increased lactate production [17–19], reduced ATP production and altered glucose utilization and transfer [16,20–22], ensures continued placental metabolism, but the energy deficit it incurs may lead to placental insufficiency, a state in which the supply of nutrients and often oxygen to the fetus is chronically suboptimal. This placental insufficiency is a major driver of IUGR and may compromise placental ATP-dependent processes, including nutrient transport, thereby contributing to adaptive fetal development and growth, which is understood to set the stage for later-life metabolic dysfunction [11,23].

While ultrasound remains the clinical standard for monitoring uteroplacental blood parameters and fetal growth, emerging imaging modalities such as hyperpolarized magnetic resonance imaging can assess placental metabolic function, a key derangement underlying IUGR [4,24,25]. Hyperpolarized [1-¹³C]pyruvate MRI enables in vivo assessment of rapid enzymatic conversion of pyruvate to downstream metabolites, including lactate and bicarbonate, providing metabolic readouts linked to LDH- and PDH-mediated routing, respectively[26][27]. These time-resolved signals can be modeled to estimate forward conversion rate constants, which reflect metabolic pathway activity within the tissue of interest [28,29]. We and others have applied hyperpolarized pyruvate MRI in pregnancy models to detect placental metabolic changes under conditions relevant to placental dysfunction, including dietary and growth-restriction paradigms[27,30]. Here, we extend this approach to a guinea pig spontaneous IUGR (spIUGR) model, hypothesizing that placental pyruvate routing will vary with IUGR severity and asymmetric (brain-sparing) growth phenotypes.

The guinea pig provides a valuable model for studying spIUGR because of its hemomonochorial placenta, relatively long gestation, precocial offspring, all which more closely resemble human pregnancy than those of other rodents [31,32]. Further, spontaneous growth restriction occurs within litters, mimicking human late-onset IUGR without the need for surgical or pharmacological intervention [33,34]. Notably, guinea pig runt fetuses show about 35% reduction in placental blood flow and nutrient transfer compared to larger littermates, mirroring altered uteroplacental blood flow and hypoxia and offering relevant context for metabolic readout studies in spIUGR [35].

In this study, we used hyperpolarized [1-¹³C]pyruvate MRI to evaluate near-term placental metabolism in a guinea pig spIUGR model and integrated these metabolic measures with structural MRI-derived fetal and placental volumes and MRI-based body composition metrics, including liver and adipose tissue proton density fat fraction (PDFF). The aims were: (1) to measure placental and fetal volumes via MRI and compare them with traditional tissue collection metrics, (2) using MRI, characterize maternal and fetal whole body and liver fat distribution and PDFF, and (3) to link placental [1-^13^C]pyruvate metabolism with IUGR outcome/severity markers. Establishing these placental metabolic signatures in a model that recapitulates key elements of human near term late-onset IUGR physiology may provide mechanistic insight into placental function under insufficiency and inform the development of more accessible, functionally targeted diagnostic strategies for earlier identification of IUGR.

## Materials and Methods

### Experimental Animals

Western University’s Animal Care Committee approved and monitored the animal care and research protocol (2021-022) in accordance with the Canadian Council on Animal Care (CCAC) standards and guidelines. Twenty-two Dunkin-Hartley guinea pig sows were acquired (Charles River Laboratories, St-Constant, Que, Canada) and allowed to acclimatize to their housing for seven days prior to monitoring for estrus for two estrus cycles prior to mating with guinea pig boars[36]. The sows had *ad libitum* access to regular guinea pig chow (LabDiet Guinea Pig Diet 5025: 60% carbohydrates, 13% fat, 26% protein; Land O’ Lakes, Inc, Arden Hills, MN) and water. Sows were individually housed in a temperature and humidity-controlled room (20°C and 30%, respectively) with a 12-hour light-dark cycle.

### 1H Imaging

Twenty-two sows were fasted for two hours prior to a magnetic resonance imaging (MRI) exam (at 60 ± 1 days gestation, full term ∼ 68 days, number of fetuses = 3.3 ± 1.0, total number of fetuses = 73) to standardize the sows’ metabolic state [30]. Sows were administered a subcutaneous injection of Robinul (glycopyrrolate, 0.01 mg/kg; Sandoz Canada, Inc., Montreal, QC, Canada) thirty minutes prior to imaging to reduce the risk of aspiration during anesthesia (induced with 4 % isoflurane, then maintained at 1.5 – 2.5 %, both with 2 L/min O_2_) [37]. The sows were warmed with a heating pad, and maternal vital signs (heart rate, respiratory rate and temperature) were monitored during imaging and recovery. All images were acquired at 3.0 Tesla (MR750, GE, Waukesha, WI, USA) using a 32-channel cardiac receive array (Invivo, Gainesville, FL, USA) for proton images and a custom-built ^13^C birdcage coil (Morris Instruments, Ottawa, Canada) for hyperpolarized images. The sows were placed in a custom animal holder for the duration of the MRI exam to minimize movement while the coils were switched between scans. Following image acquisition, sows were recovered from anesthesia and returned to their cages for at least 48 hours prior to collection. Two fetuses from a single sow with four fetuses were removed from the study as their pregnancy was aborted after imaging and prior to animal tissue collections (total number of fetuses = 71).

The MRI sequences included T_1_-weighted (T_1_-w), T_2_-weighted (T_2_-w), iterative decomposition of water and fat with echo asymmetry and least-squares estimation (IDEAL) water, fat, and proton density fat fraction (PDFF) images, multicomponent driven equilibrium single pulse observation of T_1_ and T_2_ (mcDESPOT) images and hyperpolarized [1-^13^C]pyruvate metabolic images (metabolic images) [27,30].

The 3D T_1_-w gradient-echo ^1^H images (field of view (FOV) = 20 x 0.6 cm, relaxation time/echo time (TR/TE) = 5.1/2.4 msec, flip angle = 15°, excitations = 4, slice = 0.9 mm, 0.875 x 0.875 mm^2^ in-plane resolution, ∼ 7 min acquisition time) and 3D T_2_-w spin-echo ^1^H images (FOV = 10-12 cm, TR/TE = 2002/218 msec, number of excitations = 2, excitations = 2, slice = 0.6 mm, 0.6 x 0.6 mm^2^ in-plane resolution, ∼ 10 mins acquisition time) were acquired of the whole sow for anatomical references for the hyperpolarized [1-^13^C]pyruvate metabolic images [27,30,38]. The IDEAL images (TR/TE = 9.4/0.974msec, excitations = 4, echoes = 6, flip angle = 4°, slice = 0.9 mm, 0.933 x 0.933 mm^2^ in-plane resolution, ∼ 13 mins acquisition time) were acquired of the whole sow for quantitative assessment of liver and adipose tissue depots [39,40].

### 13C Imaging

All animals underwent hyperpolarized imaging immediately after proton imaging, thirteen animals (number of fetuses with hyperpolarized data = 41) had successful hyperpolarized images acquired. Three-dimensional (3D) hyperpolarized [1-^13^C]pyruvate metabolic images were acquired using a multiphase broadband fast gradient recalled multi-echo pulse sequence (FOV = 20 x 0.6 cm, slice = 8.5 mm, bandwidth = 8.93kHz, echoes = 8, echo train length = 4, first TE = 4.2 ms, echo spacing = 1.1 ms, excitations = 1, ∼ 7.5 sec for each image, ∼1 min total acquisition time) [27,30].

The hyperpolarized 250-mM [1-^13^C]pyruvate solution consisted of [1-^13^C]pyruvic acid (Cambridge Isotope Laboratories, Tewksbury, MA) with 15-mM OX63 trityl radical (Oxford Instruments, Abingdon, United Kingdom) and 1-mM ProHance (Bracco, Milan, Italy) that was hyperpolarized with the dynamic nuclear polarizer (Hypersense, Oxford Instruments) immediately prior to injection. Once the 75 mg/kg solution was hyperpolarized, it was transported to the MRI room (∼15 seconds) and injected into the right saphenous vein of the guinea pig sow over ∼ 12 seconds. At ∼7.5 seconds after the start of the injection, imaging was initiated, and 7 total 3D images were acquired at 7.5-second intervals. At the first imaging time point, the PYR signal is observed in the hind limb following the saphenous vein injection, as imaging is started around halfway through the bolus injection. PYR is then transported to the maternal heart and then onto the fetal placenta, maternal liver, and small intestines. At these later time points, PYR, ALA, LAC and BIC signals are seen in the heart, and PYR, LAC and BIC are seen in the liver and small intestines.

### Animal Tissue Collections

The sows were euthanized 2.6 ± 0.8 days after MRI imaging (63.0 ± 1.1 days gestation) with CO_2_ inhalation [41]. The sow’s uterus was exposed, and the fetuses were confirmed dead. The locations of the fetuses were noted in a counterclockwise fashion, starting at the top of the left horn and moving down to the cervix and then back up to the top of the right uterine horn. They were removed and weighed immediately. These fetal locations from the collection were matched to imaging locations, then confirmed by ranking weight and volume from largest to smallest. The maternal liver, fetal brains, fetal livers and placentae were also weighed. The following markers of fetal growth and their respective cut-off values were used for determination of IUGR: body weight, brain-body weight ratio, brain-liver weight ratio, brain-placenta weight ratio, and body weight with respect to the average body weights in pregnancy [38,42–50]. The cut-off value for each marker was: body weight < 80 g, brain-body weight ratio > 0.03, brain-liver weight ratio > 0.7, brain-placenta weight ratio > 0.7, body weight with respect to the average body weights in pregnancy < 0.9. Each cut-off value represents the 10^th^ percentile of the population [43,47,51].

For each fetus, if the value for each spIUGR marker was above the cut-off value (brain-body weight ratio, brain-liver weight ratio, and/or brain-placenta weight ratio) or below the cut-off value (body weight and/or body weight with respect to the average body weights in pregnancy), it was noted to have one marker. Fetuses exhibiting three or more of these markers were classified as spIUGR, while those with fewer than three markers were classified as non-spIUGR.

### Segmentations

Regions of interest (ROI) were manually thresholded and segmented (Wacom Cintiq 22HD, Saitama, Japan) in 3D Slicer (version 4.13.0) on the T_1_-w, T_2_-w, IDEAL water and fat images to obtain organ volumes and median PDFF values [52]. Maternal ROI measurements included whole-body (WB), subcutaneous adipose tissue (SAT), visceral adipose tissue (VAT), and liver volumes. Placental and fetal ROI measurements included placental and total fetal volumes as well as fetal brain, liver, and adipose tissue (AT) volumes. The maternal, placental and fetal ROIs were completed by L.M. (4 years of experience). Maternal WB and fetal and maternal AT ROIs were segmented on the IDEAL water and fat images. Placental and fetal ROIs were generated from T1-w images, as they matched the hyperpolarized data. The fetal brain and liver ROIs were completed by an image analyst (6 years of experience) on T_2_-w images with higher resolution. As mentioned above, these ROIs were used for spIUGR determination with volume markers: body volume, brain-body volume ratio, brain-liver volume ratio, brain-placenta volume ratio, and body volume with respect to the average body volumes in pregnancy, where the cut-off values were calculated with linear regressions from the weight cut-off values.

The adipose tissue ROIs were eroded (maternal VAT and fetal AT: eight points once and maternal SAT: four points twice) prior to quantification to reduce partial volume effects. Maternal SAT was eroded differently to avoid the ROI disappearing with erosion. The adipose tissue ROIs were displayed as a percentage of adipose tissue volume relative to WB volume or fetal volume, respectively, to normalize for sow and fetal size effects. The ROIs were overlayed onto the PDFF images to measure the mean liver PDFF values and the median AT PDFF values. The median was used for AT PDFF values because three outliers in the maternal SAT dataset had very low values, and the median is more robust to outliers than the mean.

### Post-Processing Metabolic Images

The hyperpolarized [1-^13^C]pyruvate images of pyruvate (PYR) and each metabolite (alanine (ALA), bicarbonate (BIC), and lactate (LAC)) were reconstructed using an in-house multiphase ^13^C IDEAL reconstruction in MATLAB 2020a (MathWorks, Natick, MA) [53]. This reconstruction method used the separation of signal due to the chemical shift between PYR and its metabolites [53]. The resulting metabolic images were examined for sufficient visible signal in the placentae. The forty-one placental ROIs were overlayed onto the hyperpolarized [1-^13^C]pyruvate images to measure the mean signal intensities of [1-^13^C]pyruvate and its metabolites as a function of time. The metabolic conversion rates were estimated using the area under the curve (AUC) method, where the AUC for each metabolite (ALA, BIC and LAC) is divided by the AUC for PYR, resulting in AUC ratios (alanine-pyruvate ratio (APR), bicarbonate-pyruvate ratio (BPR) and lactate-pyruvate ratio (LPR), respectively) [29]. The spIUGR markers were correlated against the AUC ratios to observe differences in downstream metabolism.

### Statistical Analysis

Linear regressions were used to predict the spIUGR volume cut-off values from the spIUGR weight cut-off values. Paired t-tests were used to compare the number of fetuses classified as spIUGR and non-spIUGR from weight and volume measurements. A linear mixed model one-way ANOVA with a *post hoc* Tukey test with the maternal sow controlled for as a random variable, and Hedges’ g (g) indexes were calculated to assess effect size between non-IUGR and spIUGR fetal and placental weights and volumes. A small value of 0.2, a medium value of 0.5, and a large value of 0.8 are the common effect sizes for Hedges’ g [54,55]. This process was repeated for fetal adiposity measures. The fetal and maternal adiposity measures were compared using linear regression. Finally, a simple linear regression was performed between the hyperpolarized MRI metabolite ratios and two spIUGR measures: brain-body and fetal volumes and weights. The statistical tests were completed using Prism 10.2.3 (GraphPad, Boston, MA) and in RStudio (2024.04.2+764); significance was defined at p ≤ 0.05 for these tests.

## Results

### Validation of Volume Ratios Using Weight Ratios

The markers for IUGR are typically based on weight; here, we validated the volume measurements as markers for IUGR in non-invasive longitudinal growth studies. The weight cut-off value (Table 1 & Figure 1) for each spIUGR marker was used to estimate the volume cut-off value (Table 1 & Figure 2) by inputting the weight cut-off value into the linear regression equation, resulting in the estimated volume cut-off value. Six fetuses were classified as spIUGR (Table 2), and 61 fetuses were classified as non-spIUGR with both volume and weight cut-off values. The numbers of spIUGR and non-spIUGR fetuses for both volume and weight cut-off values (Table 3) were not significantly (p = 0.92 and p = 0.97, respectively) different. Six fetuses were classified as spIUGR, and sixty-one fetuses were classified as non-spIUGR with both weight and volume. Two fetuses were classified as spIUGR with only weight, and two were classified as spIUGR with only volume measurements. The number of fetuses that were classified as non-spIUGR and spIUGR for both volume and weight was consistent.

**Figure 1:**
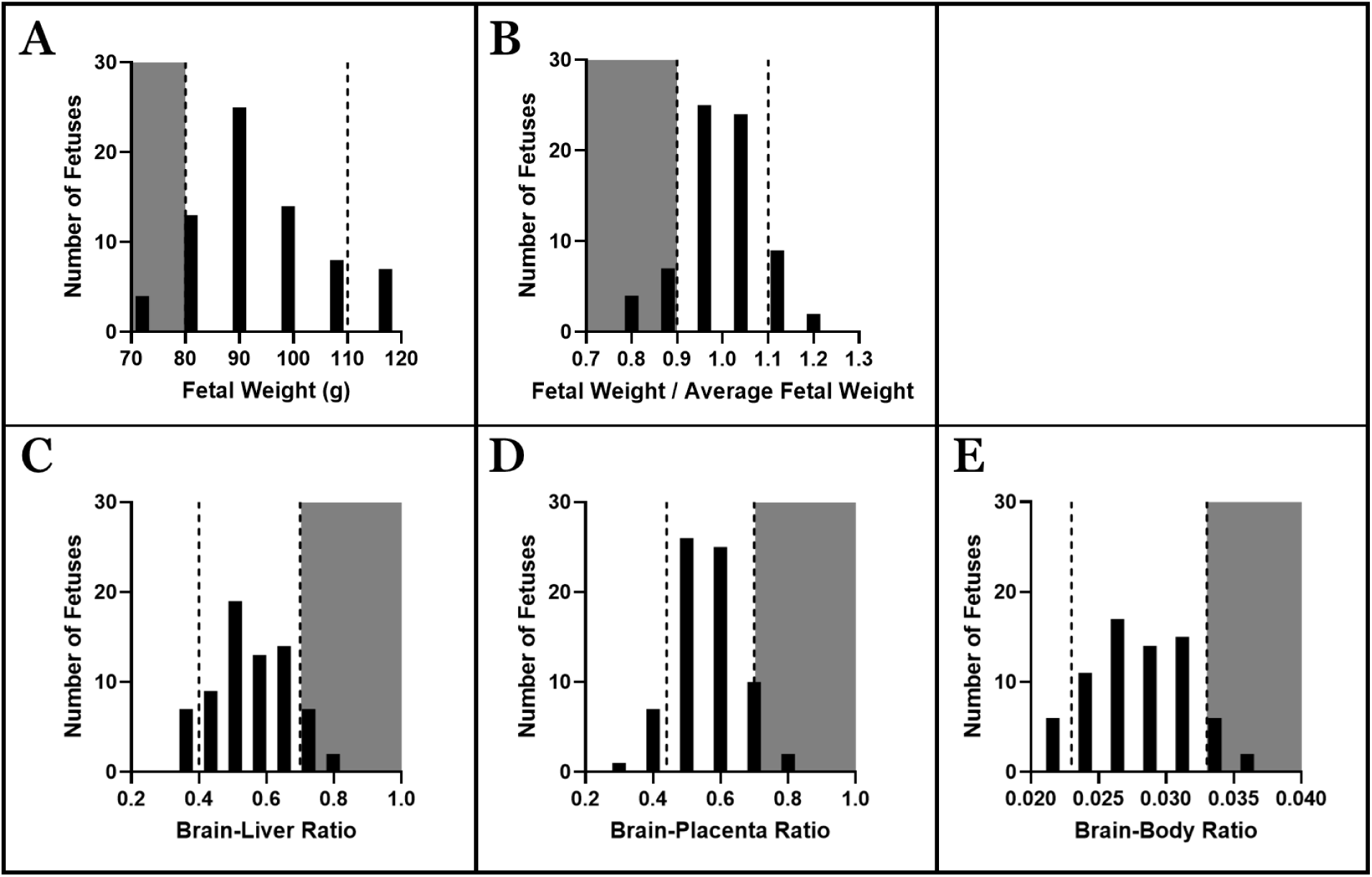
Histogram for (A) Fetal Weight (g), (B) Fetal Weight / Average Fetal Weight (fetal weight with respect to the average fetal weights in pregnancy), (C) Brain-Liver Ratio from weights, (D) Brain-Placenta Ratio from weight, and (E) Brain-Body Ratio from weight. Vertical dotted line denotes the 10th and 90th percentiles. Gray area represents the percentile that was included as the classification for spIUGR.

**Figure 2:**
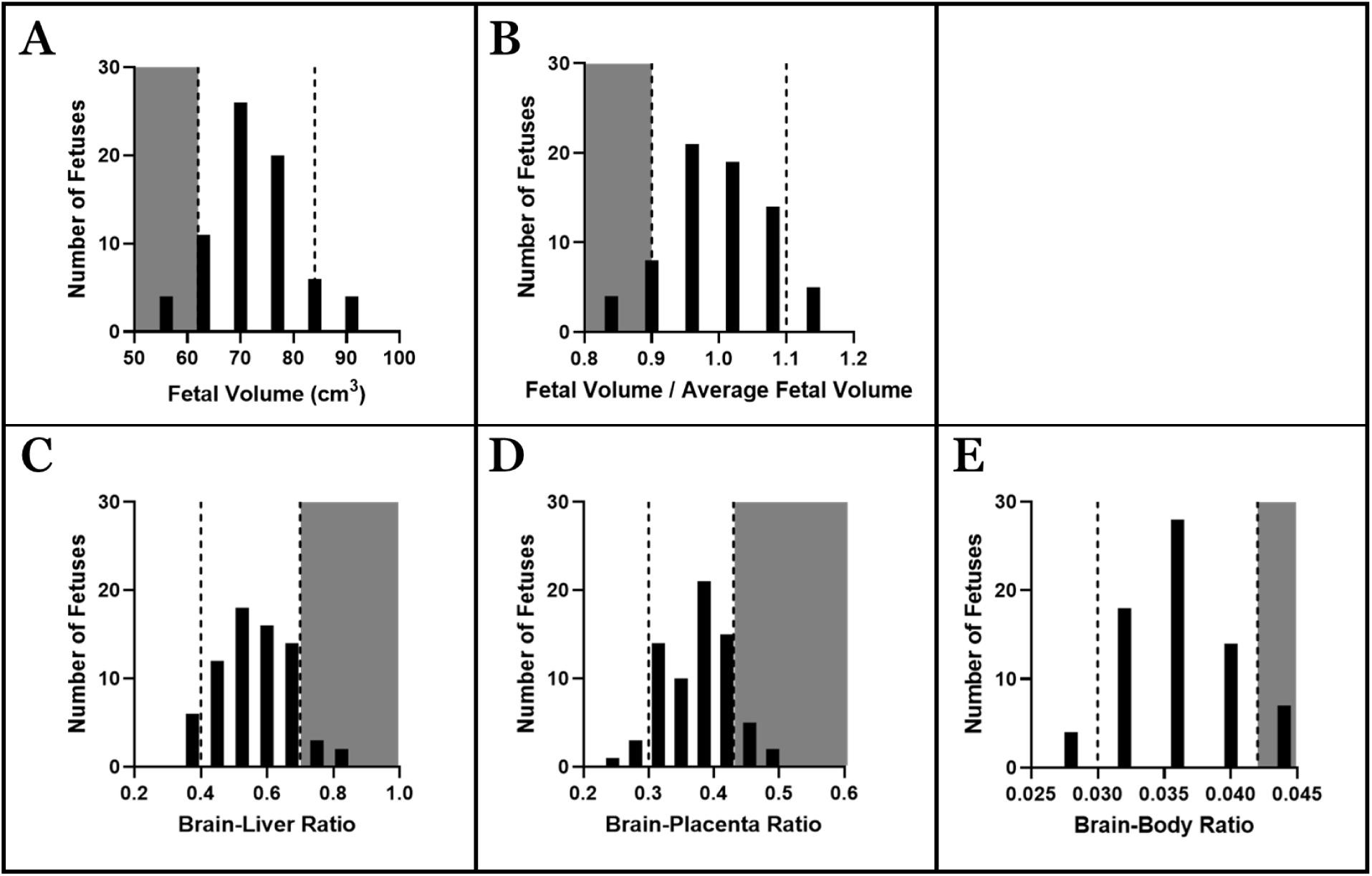
Histogram for (A) Fetal Volume (cm^3^), (B) Fetal Volume / Average Fetal Volume (fetal volume with respect to the average fetal volumes in pregnancy), (C) Brain-Liver Ratio from Volume, (D) Brain-Placenta Ratio from Volume, and (E) Brain-Body Ratio from Volume. Vertical dotted line denotes the 10th and 90th percentiles. Gray area represents the percentile that was included as the classification for spIUGR.

**Table 1:** Linear Regression Formula to Calculate spIUGR Volume cut-off values from IUGR Weight cut-off values.

| IUGR Marker | Weight Cut-Off Value | Linear Regression of Weight (g) vs. Volume (cm<sup>3</sup>) | SD | R<sup>2</sup> | Volume Cut-off Value |
| --- | --- | --- | --- | --- | --- |
| Body | 80 (g) | $Y = 0.61 * X + 15.04$ | $Y = 0.03 * X + 3.19$ | 0.83 | 64 (cm<sup>3</sup>) |
| Body with respect to the average body in pregnancy | 0.9 | $Y = 0.89 * X + 0.11$ | $Y = 0.03 * X + 0.03$ | 0.94 | 0.9 |
| Brain-body ratio | 0.03 | $Y = 1.05 * X + 0.01$ | $Y = 0.06 * X + 0.01$ | 0.84 | 0.04 |
| Brain-liver ratio | 0.7 | $Y = 0.51 * X + 0.28$ | $Y = 0.06 * X + 0.03$ | 0.53 | 0.7 |
| Brain-placenta ratio | 0.7 | $Y = 0.43 * X + 0.14$ | $Y = 0.03 * X + 0.02$ | 0.78 | 0.4 |

Footnote: spiUGR: spontaneous intrauterine growth restriction, SD: standard deviation

**Table 2:** Number of Fetuses Classified as spIUGR and non-spIUGR with each IUGR Marker using the Weight and Volume cut-off values, respectively.

| IUGR Marker | Weight spiUGR cut-off value | Non-spiUGR (n=63) | spiUGR (n=8) | Volume spiUGR cut-off value | Non-spiUGR (n=63) | spiUGR (n=8) |
| --- | --- | --- | --- | --- | --- | --- |
| Body | $\geq 80$ g | 63 | 1 | $\geq 64$ cm<sup>3</sup> | 60 | 1 |
| | $< 80$ g | 0 | 7 | $< 64$ cm<sup>3</sup> | 3 | 7 |
| Body with respect to the average body in pregnancy | $\geq 0.9$ | 61 | 2 | $\geq 0.9$ | 61 | 4 |
| | $< 0.9$ | 2 | 6 | $< 0.9$ | 3 | 3 |
| Brain-body ratio | $\geq 0.03$ | 11 | 7 | $\geq 0.04$ | 4 | 7 |
| | $< 0.03$ | 52 | 1 | $< 0.04$ | 59 | 1 |
| Brain-liver ratio | $\geq 0.7$ | 2 | 6 | $\geq 0.7$ | 4 | 2 |
| | $< 0.7$ | 61 | 2 | $< 0.7$ | 59 | 6 |
| Brain-placenta ratio | $\geq 0.7$ | 1 | 3 | $\geq 0.4$ | 18 | 7 |
| | $< 0.7$ | 62 | 5 | $< 0.4$ | 45 | 1 |

Footnote: spiUGR: spontaneous intrauterine growth restriction

**Table 3:** Differences between spIUGR Markers from Volume and Weight cut-off values.

|  |  |
| --- | --- |
| Both Volume and Weight – spIUGR | 6 |
| Only Volume – spIUGR | 2 |
| Only Weight – spIUGR | 2 |
| Both Volume and Weight – non-spIUGR | 61 |

Footnote: spIUGR: spontaneous intrauterine growth restriction

Body weight and volume with respect to the average body volume and weight in pregnancy (Figure 1B and Figure 2B), and fetal liver weight and volume (Table 4 and Table 5) were significantly lower in spIUGR fetuses than non-spIUGR fetuses. Brain weight and volume (Table 4 and Table 5) were not significantly different between non-spIUGR and spIUGR fetuses, and there was a medium and small effect size between groups for volume and weight, respectively. Similarly, placental weight and volume (Table 4 and Table 5) were not significantly different between groups, and both had large effect sizes between non-spIUGR and spIUGR fetuses. Brain-body, brain-liver, and brain-placenta weight and volume ratios were significantly higher in spIUGR fetuses than in non-spIUGR fetuses (Figure 1, Figure 2, Table 4, and Table 5).

**Table 4:** Fetal and placental weights in spIUGR.

|  | Weight |  | t ratio | g | P value |
| --- | --- | --- | --- | --- | --- |
|  | non-spIUGR | spIUGR |  |  |  |
| Body (g) | 96.4 ± 2.0 | 80.6 ± 3.9 | 4.2 | 1.4 | 0.0001 |
| Placental (g) | 4.9 ± 0.1 | 4.2 ± 0.3 | 2.0 | 0.8 | 0.05 |
| Brain (g) | 2.6 ± 0.1 | 2.5 ± 0.1 | 1.5 | 0.6 | 0.14 |
| Liver (g) | 5.0 ± 0.1 | 3.8 ± 0.3 | 3.3 | 1.3 | 0.002 |
| Body with respect to the average body in pregnancy | 1.01 ± 0.01 | 0.90 ± 0.03 | 3.8 | 1.4 | 0.0004 |
| Brain-body ratio | 0.027 ± 0.001 | 0.032 ± 0.001 | -3.7 | -1.2 | 0.0005 |
| Brain-liver ratio | 0.54 ± 0.01 | 0.69 ± 0.04 | -3.8 | -1.4 | 0.0003 |
| Brain-placenta ratio | 0.55 ± 0.01 | 0.63 ± 0.04 | -2.2 | -0.8 | 0.03 |
| Fetal-placenta ratio | 20.1 ± 0.5 | 19.6 ± 0.9 | 0.5 | 0.1 | 0.6 |

Footnote: Data are expressed as mean ± SEM for all fetuses, spIUGR: spontaneous intrauterine growth restriction

**Table 5:** Fetal and placental volumes in spIUGR.

|  | Volume |  | t ratio | g | P value |
| --- | --- | --- | --- | --- | --- |
|  | non-spIUGR | spIUGR |  |  |  |
| Body (cm<sup>3</sup>) | 73.8 ± 1.0 | 61.9 ± 2.5 | 4.8 | 2 | <0.0001 |
| Placental (cm<sup>3</sup>) | 7.1 ± 0.1 | 6.4 ± 0.3 | 1.9 | 0.7 | 0.07 |
| Brain (cm<sup>3</sup>) | 2.6 ± 0.1 | 2.6 ± 0.1 | -0.4 | -0.2 | 0.7 |
| Liver (cm<sup>3</sup>) | 4.9 ± 0.1 | 4.0 ± 0.3 | 2.3 | 1.0 | 0.03 |
| Body with respect to the average body in pregnancy | 1.01 ± 0.01 | 0.90 ± 0.02 | 4.0 | 1.5 | 0.0001 |
| Brain-body ratio | 0.035 ± 0.001 | 0.042 ± 0.001 | -6.5 | -2.4 | <0.0001 |
| Brain-liver ratio | 0.55 ± 0.01 | 0.65 ± 0.04 | -2.8 | -1.1 | 0.008 |
| Brain-placenta ratio | 0.37 ± 0.01 | 0.41 ± 0.02 | -2.5 | -1.0 | 0.013 |
| Fetal-placenta ratio | 10.5 ± 0.2 | 9.8 ± 0.4 | 1.8 | 0.7 | 0.08 |

Footnote: Data are expressed as mean ± SEM for all fetuses, spIUGR: spontaneous intrauterine growth restriction

### Body Composition Data

Maternal and fetal body composition measurements are shown in Figure 3. One maternal sow was excluded from liver analyses due to a water–fat swapping error with the iterative decomposition of water and fat with echo asymmetry and least squares estimation (IDEAL) images, and one additional sow was excluded from adipose tissue analyses.

**Figure 3:**
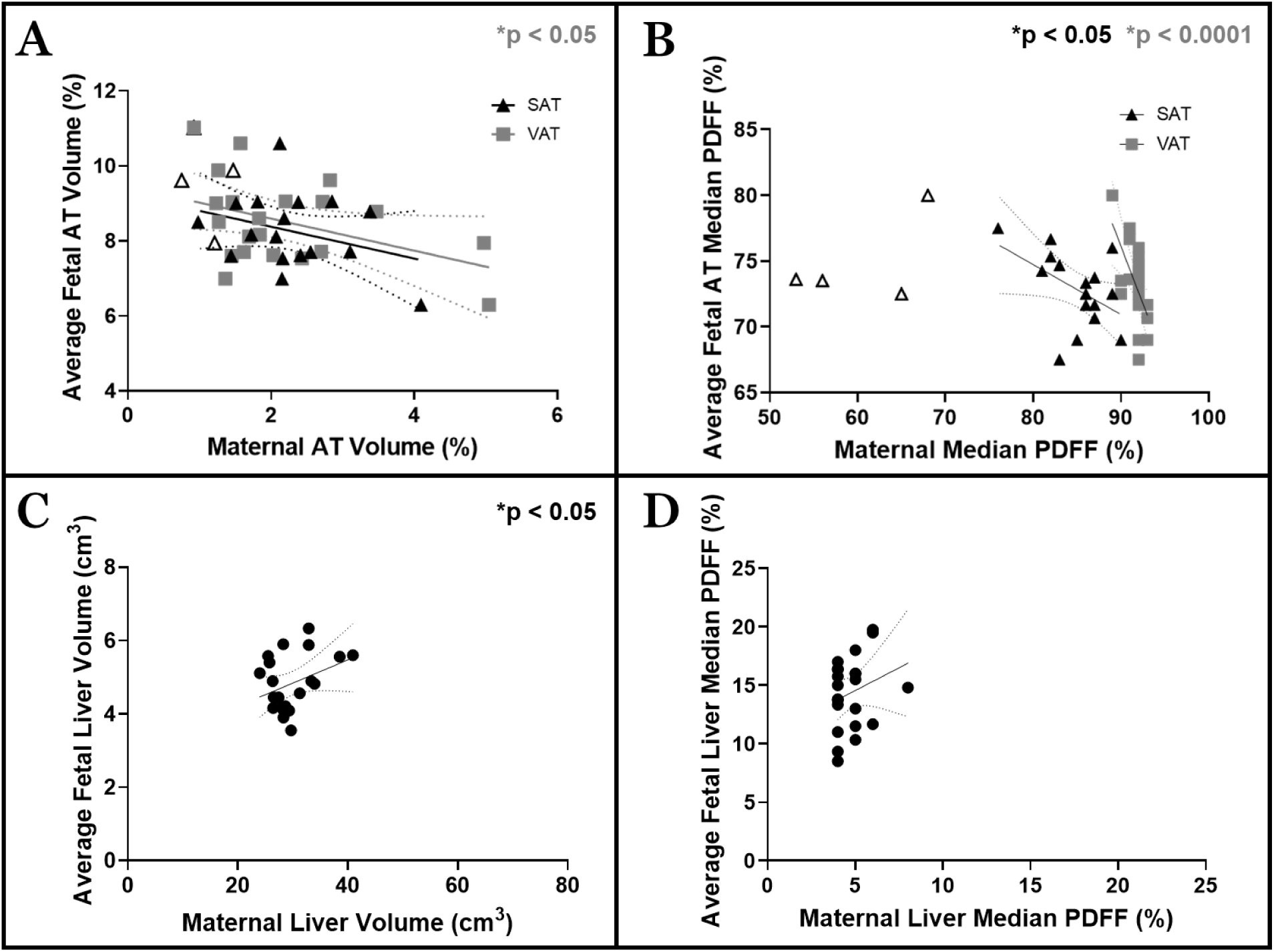
Correlation between maternal and fetal (A) adipose tissue (AT) volume percentage, (B) mean AT proton density fat fraction (PDFF), (C) liver volume (cm^3^), (D) liver mean PDFF; with linear regression lines and 95% confidence intervals. Each circle represents the average fetal measure for each maternal sow. This was done to visualize the graphs better; however, the data was analyzed with all of the fetuses. Black triangles: maternal subcutaneous AT (SAT), hollow black triangles: maternal SAT values cut from analysis, gray squares: maternal visceral AT (VAT), and black circles: liver. A percentage of AT is used to normalize the maternal and fetal size. Subcutaneous fetal AT is not observable at this time point.

Maternal adiposity was inversely associated with fetal adipose tissue (AT) development. Both maternal subcutaneous adipose tissue (SAT) and visceral adipose tissue (VAT) showed significant negative correlations with fetal AT volume (SAT; Figure 3A, F (1, 64) = 9.6, R^2^ = 0.13, slope = - 0.6, p = 0.003; VAT; Figure 3A, F (1, 64) = 5.1, R^2^ = 0.07, slope = -0.4, p = 0.03). Maternal VAT also showed a significant negative correlation with fetal AT median proton density fat fraction (PDFF; Figure 3B, F (1, 64) = 25.6, R^2^ = 0.29, slope = -1.6, p = <0.0001), while the relationship between maternal SAT and fetal AT PDFF was not significant (Figure 3B, F (1, 64) = 3.5, R^2^ = 0.05, slope = -0.1, p = 0.07).

Four sows displayed notably low SAT PDFF values (Figure 3B). These outliers were verified as low SAT volume, reflecting low lipid accumulation, and were not reliable measurements due to partial volume effects. A reanalysis excluding these animals strengthened the maternal SAT–fetal AT PDFF relationship to significance (Figure 3B, F(1,49) = 5.4, R² = 0.10, slope = −0.3, p = 0.02) but the maternal SAT–fetal AT volume relationship lost significance (Figure 3A, F(1,49) = 0.4, R² = 0.01, slope = −0.2, p = 0.5). Finally, Fetal AT volume (%) and median PDFF did not differ significantly between spIUGR and non-spIUGR fetuses (Table 6).

**Table 6:** Fetal adiposity in spIUGR on fetal adiposit.

|  | non-spIUGR | spIUGR | t ratio | g | P value |
| --- | --- | --- | --- | --- | --- |
| Fetal Liver Volume (cm<sup>3</sup>) | 4.9 ± 0.1 | 4.0 ± 0.3 | 2.3 | 1.0 | 0.03 |
| Fetal Liver Median PDFF | 14.5 ± 0.7 | 14.5 ± 1.0 | -0.1 | 0.1 | 0.99 |
| Fetal AT volume (%) | 0.09 ± 0.01 | 0.08 ± 0.01 | 1.5 | 0.1 | 0.14 |
| Fetal AT Median PDFF | 73.3 ± 0.6 | 74.0 ± 0.8 | -1.1 | -0.3 | 0.3 |

Footnote: Data are expressed as mean ± SEM for all fetuses, AT: Adipose Tissue, spIUGR: spontaneous intrauterine growth restriction, PDFF: proton density fat fraction

For liver measurements, maternal and fetal liver volumes were positively correlated (Figure 3C, F (1, 69) = 4.8, R^2^ = 0.06, slope = 0.1, p = 0.032), though the correlation between maternal and fetal liver median PDFF did not reach significance (Figure 3D; F(1,69) = 3.4, R² = 0.05, slope = 0.3, p = 0.07). Fetal liver volume was significantly lower in spIUGR compared to non-spIUGR fetuses (Table 6), while liver median PDFF did not differ between groups.

### Fetoplacental Metabolism

Hyperpolarized [1-^13^C]pyruvate MRI demonstrated phenotype-dependent differences in downstream metabolite conversion as shown in Figure 4. Linear regressions between hyperpolarized MRI lactate and bicarbonate pyruvate ratios (LPR and BPR, respectively) and all spIUGR markers for weight (body weight, brain-body weight ratio, brain-liver weight ratio, brain-placenta weight ratio, body weight with respect to the average body weights in pregnancy) and volume were generated (Supplemental Data: Table S1 and Table S2). In our cohort, brain-body weight/volume ratios (Figure 5) were elevated across for LPR, and body weight/volume were reduced for BPR across both weight and volume measurements. The other spIUGR markers did not show significant correlation for both weight and volume measurements (Supplemental Data: Figure S1, Figure S2, Figure S3, Figure S4).

**Figure 4:**
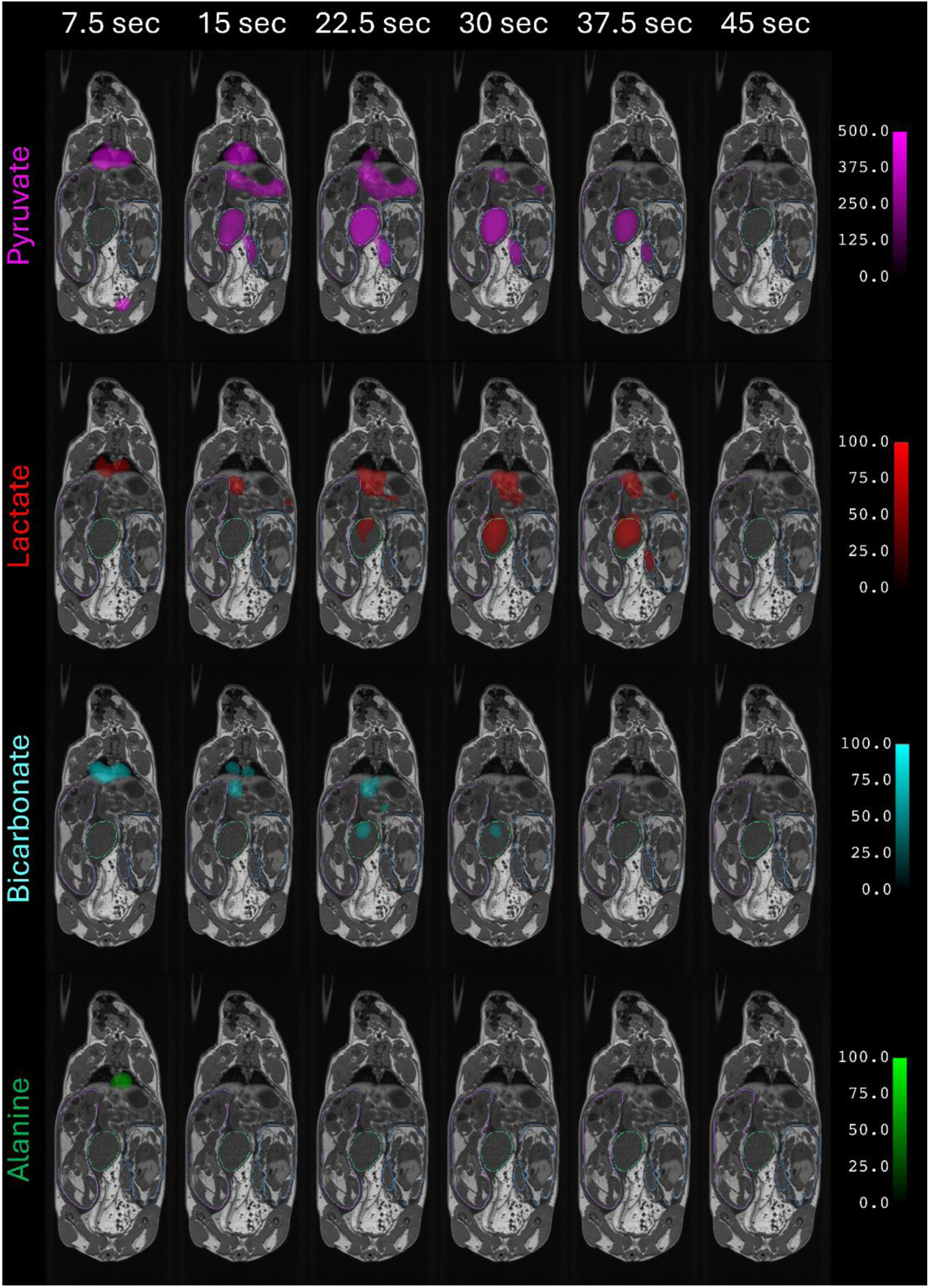
A 61-day pregnant sow with three fetuses. T_1_-weighted image with first 6 of 7 timepoints of metabolic images (pyruvate: magenta, lactate: red, bicarbonate: cyan, alanine: green) overlayed to observe signal intensity over time. In this image slice, one fetus (far left, contoured in purple), one placenta (middle, contoured in green), one fetus and matching placenta are visible (far right, contoured in blue). Alanine has no visible signal in the placenta. The pyruvate images and the metabolite images are windowed and levelled identically (Min: 0, Max: 500, Lower Threshold: 250 & Min: 0, Max: 100, Lower Threshold: 50, respectively).

**Figure 5:**
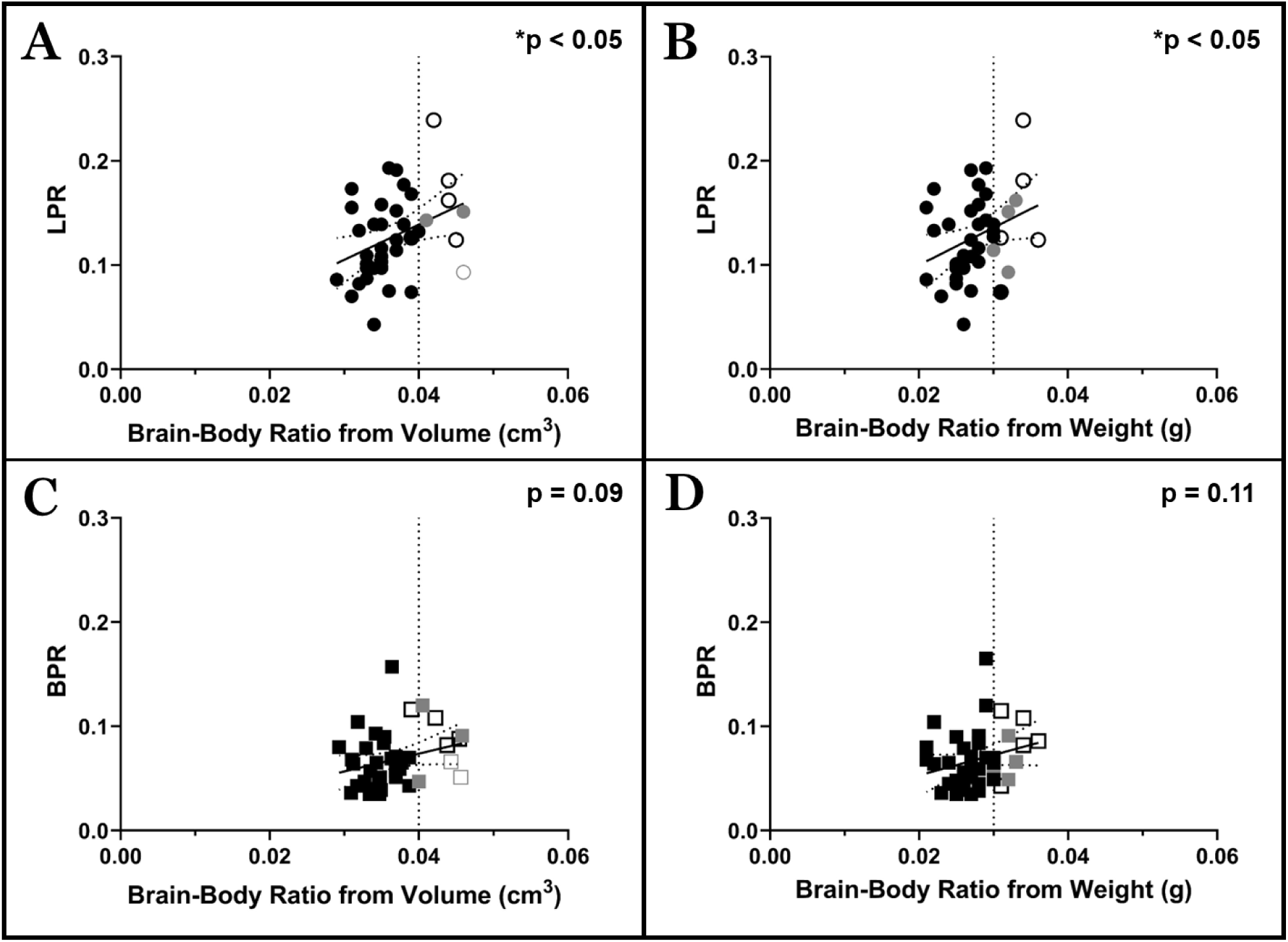
Linear regression for (A) lactate pyruvate ratio (LPR: circles) and brain-body volume (cm^3^) ratio, (B) LPR and brain-body weight (g) ratio, (C) bicarbonate pyruvate ratio (BPR: squares) and brain-body volume ratio, and (D) BPR and brain-body weight ratio, with 95% confidence intervals. The dotted line for each represents the cut-off value for the spontaneous intrauterine growth restricted (spIUGR) marker (0.04 cm^3^ or 0.03 g). Full black colour: non-spIUGR fetuses, outline black colour: spIUGR fetuses (three or more markers for spIUGR), and full gray: non-spIUGR fetuses that had that marker as spIUGR. The metabolite alanine (alanine pyruvate ratio (APR)) was not included due to low signal-to-noise in the placenta.

Brain-body weight and volume ratios (Figure 5) reflect the clinically translatable brain-sparing (asymmetric growth) adaptation to placental insufficiency [45,48,50,56]. Lactate-to-pyruvate ratio (LPR) increased with asymmetric growth, showing significant positive associations with both brain-to-body volume ratio (Figure 5A, F = 5.9 (1, 39), R^2^ = 0.13, slope = 3.38 ± 1.39, p = 0.02) and brain-body weight ratio (Figure 5B, F (1,39) = 4.5, R^2^ = 0.10, slope = 3.59 ± 1.69, p = 0.04). In contrast, bicarbonate-to-pyruvate ratio (BPR) was not significantly related to brain-to-body ratios (Figure 5C, F (1,39) = 3.1, R^2^ = 0.07, slope = 1.71 ± 0.98, p = 0.09) or brain-body weight ratio (Figure 5D, F (1,39) = 2.7, R^2^ =0.06, slope = 1.96 ± 1.18, p = 0.11).

When examined against absolute fetal size (Figure 6), LPR was significantly inversely related to fetal body volume (Figure 6A, F (1, 39) = 6.3, R^2^ = 0.14, slope = -0.0018 ± 0.0007, p = 0.02), with a similar trend for body weight (Figure 6B, F (1, 39) = 3.0, R^2^ = 0.07, slope = -0.0008 ± 0.0005, p = 0.09). BPR showed significant negative associations with both fetal body volume (Figure 6C, F (1,39) = 7.1, R^2^ = 0.15, slope = -0.0013 ± 0.0005, p = 0.02) and fetal weight (Figure 6D, F (1,39) = 6.1, R^2^ = 0.14, slope = -0.0008 ± 0.0003, p = 0.02).

**Figure 6:**
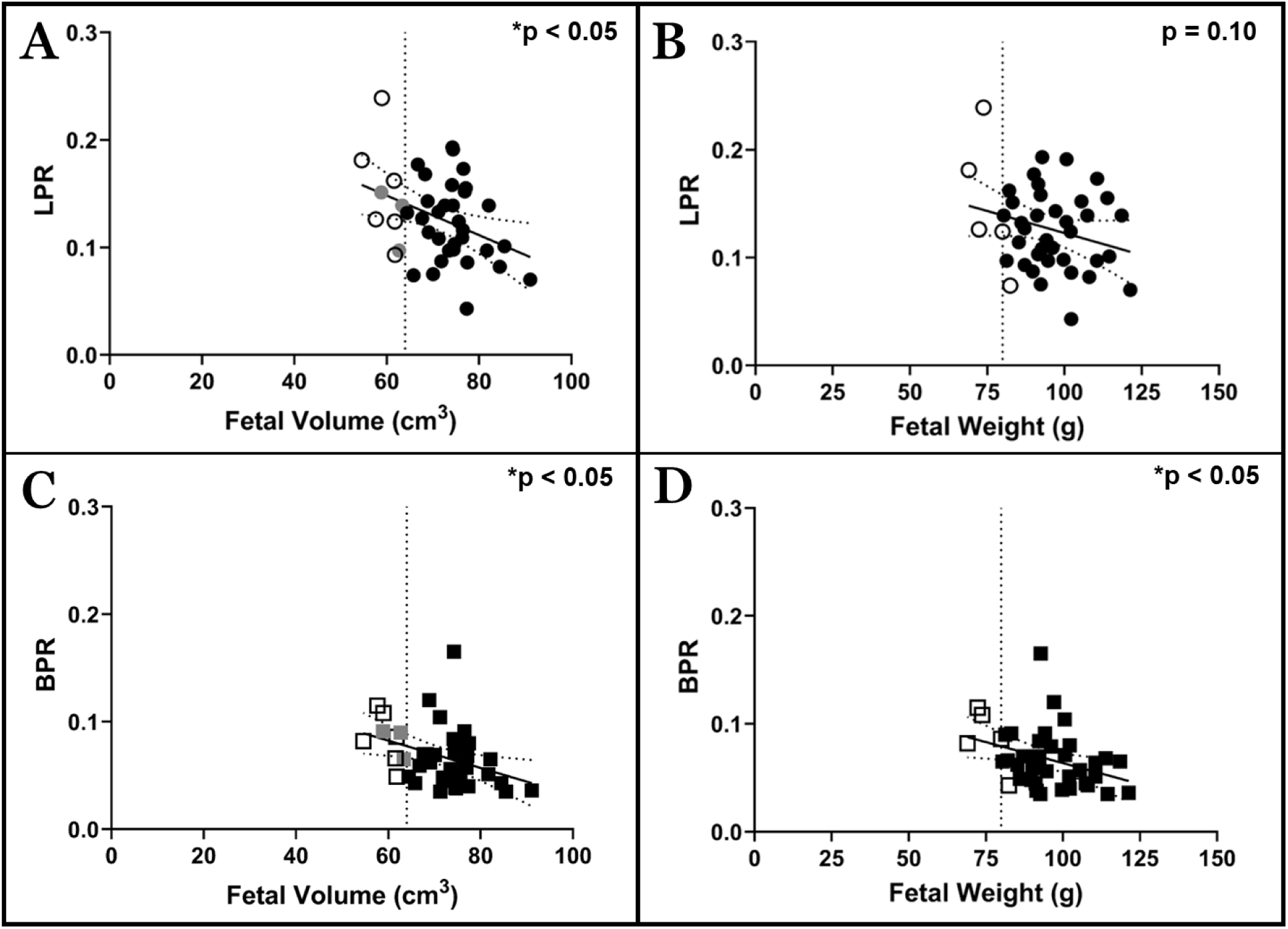
Linear regression for (A) lactate pyruvate ratio (LPR: circles) and fetal volume (cm^3^), (B) LPR and fetal weight (g), (C) bicarbonate pyruvate ratio (BPR: squares) and fetal weight (g), (D) BPR and fetal volume (cm^3^), with 95% confidence intervals. The dotted line for each represents the cut-off value for the spontaneous intrauterine growth restricted (spIUGR) marker (67 cm^3^ or 85 g). Full black colour: non-spIUGR fetuses, outline black colour: spIUGR fetuses (three or more markers for spIUGR), and full gray: non-spIUGR fetuses that had that marker as spIUGR. The metabolite alanine (alanine pyruvate ratio (APR)) was not included due to low signal-to-noise in the placenta.

## Discussion

This study measured placental and fetal volumes using MRI, compared them with weight measures, determined maternal and fetal body composition characteristics and correlated placental [1-^13^C]pyruvate metabolism with IUGR markers in a placental insufficiency spIUGR guinea pig model near term. To accomplish this, MRI images were segmented to obtain volumes, which were validated against weight measurements collected at autopsy. Using these validated volumes, this study then investigated the association of maternal metabolic health, as determined by body composition with placental metabolism and fetal body composition measurements in a spIUGR model. The body composition of the maternal and fetal volumes was measured with water, fat, and proton density fat fraction (PDFF) images. The placental volumes were overlaid on the metabolic images to observe the metabolism of pyruvate to each of its metabolites in the placenta. Our results demonstrate that the measured MRI volumes can provide the same information as organ weights, consistently identifying spIUGR. Further, this study identified altered placental metabolism in the spIUGR placenta, suggesting that this is a response to maintain placental energy levels in support of fetoplacental metabolism.

### spIUGR Markers

Spontaneous IUGR is associated with reduced placental and fetal weight in larger litter sizes, as well as placental structural and functional alterations [34,57–59]. The measured volumes correlated strongly with the weight measurements; supporting the validity of MRI generated volumes to identify spIUGR fetuses non-invasively, a key aspect of clinical medicine. Fetal, placental, and fetal liver volume/weight were decreased in spIUGR fetuses compared to non-spIUGR fetuses, consistent with previous weight studies [34,60–62]. We found that the brain volume/weight was similar between spIUGR and non-spIUGR fetuses, consistent with other reports [33,62]. The brain-liver ratio was increased in spIUGR fetuses relative to non-spIUGR fetuses, consistent with previous studies with asymmetrical spIUGR [34,60–63]. Finally, spIUGR fetuses had a significantly higher brain-body ratio than non-spIUGR fetuses, due to a lower body volume in spIUGR fetuses, while brain volume remained the same. The preserved brain size with reduced body and liver size and elevated brain-based ratios is consistent with an asymmetric growth pattern that resembles common clinical presentations of late-onset IUGR, where growth deviation can be modest on standard biometry yet redistribution is evident [6,7]. Finally, body weight-placenta weight ratio, more commonly known as placental efficiency, was measured in this study but was not used as a marker of spIUGR for this project because, in a polytocous species such as guinea pigs, placental efficiency can vary with litter size [57,64]. Additionally, using body-placenta weight (and volume) ratio as a marker for the nutrient transfer capacity of the placenta to meet the needs of the fetus is less conclusive in human studies than it is in mouse studies [65].

### Body Composition Data

Fetal adipose tissue (AT) volume (%) was reduced and negatively associated with both maternal subcutaneous AT (SAT) and visceral AT (VAT) volume (%), shown in Figure 3. Additionally, fetal AT PDFF was also negatively associated with maternal VAT PDFF but not with maternal SAT PDFF. We hypothesized that there would be an association between maternal and fetal AT depots, as previous research has shown that maternal adiposity may affect placental function and alter fetal adipose tissue development [66–69]. The negative association between fetal AT and maternal VAT PDFF may be due to the interscapular location of most of the fetal AT, a known location for brown adipose tissue (BAT) that has a lower PDFF than white adipose tissue (WAT) [70,71]. The lack of an association between fetal AT PDFF and maternal SAT PDFF is likely due to the four outlier maternal sows (Figure 3) that had very low SAT PDFF values. Due to the lower maternal SAT volume, there are more partial volume effects in the PDFF measurement.

The lower fetal liver and adipose tissue volumes observed in spIUGR fetuses relative to non-IUGR fetuses and the similar brain volume between the two groups further support development of asymmetrical spIUGR [9], and the lower adipose tissue volume in spIUGR fetuses is consistent with human studies [72]. In IUGR pregnancies, impaired placental perfusion and nutrient transport limit fetal growth despite maternal stores [73]. Our results align with the concept of asymmetric IUGR due to uteroplacental insufficiency, wherein fetal fat and nutrient storage are sacrificed to prioritize vital organs (brain-sparing effect).

### Metabolic Data

In the presence of oxygen, pyruvate can be converted into acetyl-CoA, releasing carbon dioxide that is in rapid equilibrium with bicarbonate (BIC) or oxaloacetate to fuel the TCA cycle, alanine (ALA) for protein synthesis and lipogenesis, and lactate (LAC) for energy (adenosine triphosphate (ATP). When oxygen is limited, pyruvate is converted into LAC via anaerobic glycolysis, reducing the acetyl-CoA production [74,75]; similarly, when ATP is limited, regardless of the presence of oxygen, the cell may utilize most of the pyruvate to generate LAC [76,77]. The metabolic switch from oxidative phosphorylation to aerobic glycolysis is a hallmark of cellular metabolic dysfunction [78]. Though aerobic glycolysis is 100 times faster than oxidative phosphorylation, aerobic glycolysis is a less efficient method of generating ATP, often referred to as the Warburg effect and occurs in proliferating tissues, such as tumours, but also the placenta [76–78].

Metabolic images from hyperpolarized [1-^13^C]pyruvate MRI were used to estimate the forward rate constants for conversion of pyruvate to each of its metabolites in the placentae, lactate (k_PL_), bicarbonate (k_PB_), and alanine (k_PA_). These conversion rates are directly proportional to the concentrations of the enzymes responsible for these metabolic conversions [79]. The enzymes lactate dehydrogenase (LDH) and alanine aminotransferase (ALT) are responsible for catalyzing the reaction between pyruvate and LAC and pyruvate and ALA, respectively. Pyruvate is also transported into the mitochondria, and pyruvate dehydrogenase (PDH) catalyzes the conversion from pyruvate to acetyl-CoA, releasing water and ^13^CO_2_ (BIC with carbonic anhydrase (CA)) as a part of the BIC buffer system to maintain pH levels. We observed changes in the conversion rates for K_PL_ and k_PB_ in a spIUGR model.

Hyperpolarized [1-¹³C]pyruvate MRI revealed phenotype-dependent placental substrate converison. The lactate:pyruvate ratio (LPR) showed a positive correlation with both the brain-body volume ratio and brain-body weight ratio, indicating that a more “brain-sparing” (asymmetric) growth phenotype is associated with increased hyperpolarized pyruvate-to-lactate conversion (k_PL_). This would be consistent with a hypoxia-adaptive shift toward glycolytic support within the fetoplacental unit, where LDH-mediated reduction of pyruvate to lactate regenerates NAD⁺, enabling continued glycolytic ATP production when oxidative metabolism is constrained. This is supported by primary trophoblast and trophoblast-like models, where hypoxic exposure increases lactate production alongside increased LDH expression/activity, supporting the biological plausibility of higher LPR under reduced oxygen availability [17]. More recent uteroplacental flux work further shows that placental insufficiency alters fetal nutrient handling and is associated with increased fetal lactate in PI-IUGR, highlighting lactate as a key component of the adaptive, and potentially maladaptive, metabolic response [23]. Finally, recent human data support the concept that hypo-perfused IUGR placentae exhibit metabolic reprogramming toward glycolytic pathways, aligning with the interpretation that increased LPR in spIUGR reflects sustained reliance on glycolytic metabolism under limited oxygen delivery [11]. Mechanistically, hypoxia-inducible factor 1α (HIF-1α) is a central regulator orchestrating trophoblast hypoxic adaptation. Under low oxygen, HIF-1α becomes stabilized and promotes glycolytic capacity by upregulating glycolytic programs, including LDHA and glucose transport pathways, thereby supporting pyruvate-to-lactate conversion readouts observed in our data [14,15,80]. Although our HP MRI readouts do not directly quantify fetal metabolism, independent in vivo and clinical studies support coupled fetoplacental substrate handling during hypoxia/placental insufficiency. Large-animal hypoxemia studies report altered lactate–pyruvate exchange across the placenta [81], and human preterm and term IUGR cord-blood profiles show increased placental lactate provision to the fetus with reduced fetal lactate production [11]. These observations provide a physiological framework consistent with our finding of increased placental LPR in more severely growth-restricted and brain-sparing nearterm phenotypes, while recognizing that the present measurements report placental metabolite conversions rather than fetal fluxes per se. Of note, if similar metabolic routing occurs in human pregnancies, elevated placental LPR could provide a functionally targeted signature of placental insufficiency in scenarios where late-onset IUGR may be underdetected by size metrics or umbilical artery Doppler alone.

In parallel, the bicarbonate:pyruvate ratio (BPR) was also significantly increased in smaller fetuses, with a similar (non-significant) trend toward higher BPR in those associated with brain-sparing. BPR is an index of pyruvate flux through pyruvate dehydrogenase into the TCA cycle, yielding CO₂ (detected as bicarbonate) and thus an elevated BPR points to oxidative metabolism alongside glycolysis. Notably, the concurrent rise in both placental BPR and LPR suggests an overall high-turnover metabolic state in which two arms of pyruvate fate (LDH and PDH) are upregulated. Interestingly, increased placental PDH activity has also been reported in placental insufficiency-induced IUGR placentae, though in contrast, a maternal hypoxemia study found no differences in PDH activity between hypoxic and control placentae [23,81]. While HIF-1α signaling is often linked to reduced reliance on oxygen-intensive metabolism (e.g., via induction of PDK1) [14,82], hypoxia adaptation in the placenta can be heterogeneous and compartment-dependent, and oxidative routing of a fraction of pyruvate may be preserved in the IUGR placenta [83,84]. In this framework, elevated BPR alongside LPR is compatible with coordinated placental metabolic remodeling under hypoxia, enhanced lactate exchange together with maintained or variable oxidative pyruvate routing. The rise in placental BPR alongside LPR underscores a coordinated metabolic adaptation in which the IUGR fetus–placenta unit channels pyruvate into both lactate (anaerobic route) and CO₂/bicarbonate (aerobic route), consistent with HIF-1–mediated reprogramming that supports oxygen sparing and substrate flexibility. In this framework, the placenta shifts toward glycolysis to reduce its own oxygen demand, while the brain-sparing fetus may preserve oxidative metabolism in preferentially perfused organs. External pregnancy hypoxia models (including high altitude) support an oxygen-sparing placental strategy in which placental metabolism adapts to promote fetal oxygen delivery [80], providing additional context for interpreting the LPR/BPR pattern as a placental metabolic phenotype linked to growth restriction severity. Taken together, the LPR and BPR findings suggest that growth restriction severity and growth asymmetry capture distinct but overlapping aspects of fetoplacental metabolic adaptation, with brain-sparing preferentially associated with increased placental pyruvate–lactate exchange, and reduced fetal size associated with globally increased pyruvate throughput across both glycolytic and oxidative pathways.

Finally, while ample PYR, BIC and LAC signals were found in the placentae of the metabolic images (Figure 4) for analysis, there was insufficient signal found for ALA, and hence ALA analysis was not undertaken. The low ALA signal observed in the placenta is consistent with previous results from our laboratory in other model settings [30]. Similar studies in chinchilla and rat pregnancy models also observed minimal to no ALA production in the placenta [85,86]. The low ALA signal suggests that a negligible amount of ALA is being produced in the placentae.

Decreased ALA production may be due to an increased use of LAC for energy instead of ALA and has been found in the hypoxic fetoplacental unit [81].

### Limitations and Future Work

Guinea pigs are a suitable animal model for studies of pregnancy in many ways, given important similarities with human pregnancy, including the type of placenta, a relatively long pregnancy, smaller litter sizes, and the development of fat in utero; however, there are some key differences between human and guinea pig pregnancies [87,88]. One difference between humans and guinea pigs is the distribution of adipose tissue, where humans have fetal SAT to visualize in MRI, and guinea pig fetuses do not [89]. Additionally, it is also important to note that there is significantly elevated triglyceride content in the guinea pig fetal liver, resulting in an increased liver PDFF, which is not seen in human fetal livers [90–92]. These factors need to be considered when interpreting results; however, relative changes are important and can provide translational information for pregnancy.

This study was cross-sectional to allow validation of the MRI volumes with the collection weights. Undertaking a longitudinal study would allow for the birth of the fetuses and observation of neonatal catch-up growth and potential later development of metabolic disease [34,93,94]. As the MRI volumes were validated in this study, future work could utilise these techniques and, in longitudinal studies, observe an earlier time point for identifying spIUGR fetuses and associated placental metabolic function.

Finally, in this study, simple linear regression was used to correlate the data. A multiple linear regression model including sex could further our understanding of adaptations of placental metabolism and body composition in spIUGR fetuses. However, a larger dataset would have been needed to use a multiple linear regression model, which would have been a more appropriate statistical test for analyzing the hyperpolarized data relative to multiple spIUGR measures.

## Conclusion

This study validated MRI-derived fetal and placental volume measurements against autopsy weights in a near-term placental insufficiency associated spontaneous IUGR (spIUGR) guinea pig model, supporting MRI volumetry as a non-invasive approach for identifying spIUGR in longitudinal pregnancy studies. Volume measurements reproduced spIUGR markers consistent with asymmetric growth (reduced fetal/placental/liver size with preserved brain volume and increased brain:liver and brain:body ratios), and body composition imaging showed reduced fetal adiposity with inverse relationships to maternal adiposity, particularly visceral fat metrics, highlighting reduced fetal lipid accretion in spIUGR. Using hyperpolarized [1-¹³C]pyruvate MRI, we further demonstrated phenotype-dependent placental pyruvate metabolism, with increased pyruvate-to-lactate conversion associated with more brain-sparing growth and increased pyruvate-to-bicarbonate conversion in smaller fetuses, consistent with altered placental metabolic routing in growth restriction. Together, these findings link non-invasive growth and composition markers to placental metabolic signatures and motivate longitudinal studies to determine whether these metabolic patterns precede measurable growth deviation and can inform clinically translatable diagnostic strategies.

## Acknowledgements

The authors thank Simran Sethi for their assistance with the segmentation of the fetal brain and fetal liver images. The authors acknowledge the Anishinaabek, Haudenosaunee, Lūnaapéewak and Chonnonton Nations, whose traditional territories are where this work was produced.

## Author contributions

**Lindsay E. Morris:** investigation, review & editing, visualization, writing – original draft, **Lanette J. Friesen-Waldner:** investigation, provision, project administration, writing – review & editing, **Trevor P Wade:** investigation, software, writing – review & editing, **Barbra de Vrijer**: conceptualization, writing – review & editing, supervision, funding acquisition, **Timothy R.H. Regnault:** conceptualization, writing – review & editing, supervision, funding acquisition, **Charles A McKenzie:** conceptualization, writing – review & editing, supervision, funding acquisition.

## Abbreviations

AUC: area-under-the-curve
BPR: bicarbonate pyruvate ratio
LPR: lactate pyruvate ratio
MRI: magnetic resonance imaging
spIUGR: spontaneous intrauterine growth restriction

## Footnotes

1 AUC: area-under-the-curve, BPR: bicarbonate pyruvate ratio, IUGR: intrauterine growth restriction, LPR: lactate pyruvate ratio, MRI: magnetic resonance imaging, spIUGR: spontaneous intrauterine growth restriction.

