## Supplemental Data for "Altered Placental [1-^13^C]Pyruvate Metabolism in Spontaneous Intrauterine Growth Restricted near-term Pregnant Guinea Pigs"

**Supplementary Data**

**Table S1: Lactate pyruvate ratio and spontaneous intrauterine growth restricted (spIUGR) markers with volume ($cm^3$) weight (g) measurements**

| | LPR – Volume ($cm^3$) | | | | |
|---|---|---|---|---|---|
| | Slope | Y-intercept | F | $R^2$ | p value |
| Body volume ($cm^3$) | -0.0018 ± 0.0007 | 0.26 ± 0.05 | 6.3 | 0.14 | 0.02 |
| Body volume with respect to the average body in pregnancy | -0.02 ± 0.05 | 0.09 ± 0.05 | 0.1 | 0.01 | 0.71 |
| Brain-body ratio from volume | 3.38 ± 1.39 | 0.01 ± 0.05 | 5.9 | 0.13 | 0.02 |
| Brain-liver ratio from volume | 0.01 ± 0.06 | 0.12 ± 0.04 | 0.01 | 0.01 | 0.91 |
| Brain-placenta ratio from volume | -0.01 ± 0.13 | 0.13 ± 0.05 | 0.1 | 0.01 | 0.99 |
| Fetal-placenta ratio from volume | -0.01 ± 0.01 | 0.24 ± 0.05 | 5.6 | 0.12 | 0.02 |
| | LPR – Weight (g) | | | | |
| | Slope | Y-intercept | F | $R^2$ | p value |
| Body weight (g) | -0.0008 ± 0.0005 | 0.20 ± 0.05 | 2.8 | 0.1 | 0.10 |
| Body weight with respect to the average body in pregnancy | -0.03 ± 0.07 | 0.16 ± 0.07 | 0.2 | 0.01 | 0.66 |
| Brain-body ratio from weight | 3.59 ± 1.69 | 0.03 ± 0.05 | 4.5 | 0.1 | 0.04 |
| Brain-liver ratio from weight | 0.02 ± 0.06 | 0.12 ± 0.03 | 0.07 | 0.01 | 0.79 |
| Brain-placenta ratio from weight | 0.04 ± 0.06 | 0.11 ± 0.03 | 0.35 | 0.01 | 0.55 |
| Fetal-placenta ratio from weight | -0.01 ± 0.01 | 0.16 ± 0.05 | 0.5 | 0.01 | 0.47 |

Footnote: Data are expressed as mean ± SEM for all fetuses, LPR: lactate pyruvate ratio

**Table S2: Bicarbonate pyruvate ratio and spontaneous intrauterine growth restricted (spIUGR) markers with volume (cm3) weight (g) measurements**

| | BPR – Volume ($cm^3$) | | | | |
|---|---|---|---|---|---|
| | Slope | Y-intercept | F | $R^2$ | p value |
| Body volume ($cm^3$) | -0.0013 ± 0.0005 | 0.16 ± 0.04 | 6.4 | 0.14 | 0.02 |
| Body volume with respect to the average body in pregnancy | -0.02 ± 0.1 | 0.15 ± 0.08 | 0.1 | 0.01 | 0.76 |
| Brain-body ratio from volume | 1.71 ± 0.98 | 0.01 ±0.04 | 3.1 | 0.07 | 0.09 |
| Brain-liver ratio from volume | 0.04 ± 0.04 | 0.04 ± 0.02 | 0.9 | 0.02 | 0.36 |
| Brain-placenta ratio from volume | 0.01 ± 0.08 | 0.06 ± 0.03 | 0.02 | 0.01 | 0.90 |
| Fetal-placenta ratio from volume | -0.01 ± 0.01 | 0.12 ± 0. 04 | 1.9 | 0.05 | 0.17 |
| | BPR – Weight (g) | | | | |
| | Slope | Y-intercept | F | $R^2$ | p value |
| Body weight (g) | -0.0008 ± 0.0003 | 0.14 ± 0.03 | 5.6 | 0.13 | 0.02 |
| Body weight with respect to the average body in pregnancy | -0.02 ± 0.05 | 0.08 ± 0.05 | 0.1 | 0.01 | 0.74 |
| Brain-body ratio from weight | 1.96 ± 1.18 | 0.01 ± 0.03 | 2.7 | 0.06 | 0.11 |
| Brain-liver ratio from weight | 0.04 ± 0.04 | 0.04 ± 0.02 | 1.0 | 0.02 | 0.31 |
| Brain-placenta ratio from weight | 0.05 ± 0.04 | 0.04 ± 0.02 | 1.2 | 0.03 | 0.28 |
| Fetal-placenta ratio from weight | -0.01 ± 0.01 | 0.07 ± 0.03 | 0.03 | 0.01 | 0.85 |

Footnote: Data are expressed as mean ± SEM for all fetuses, BPR: bicarbonate pyruvate ratio

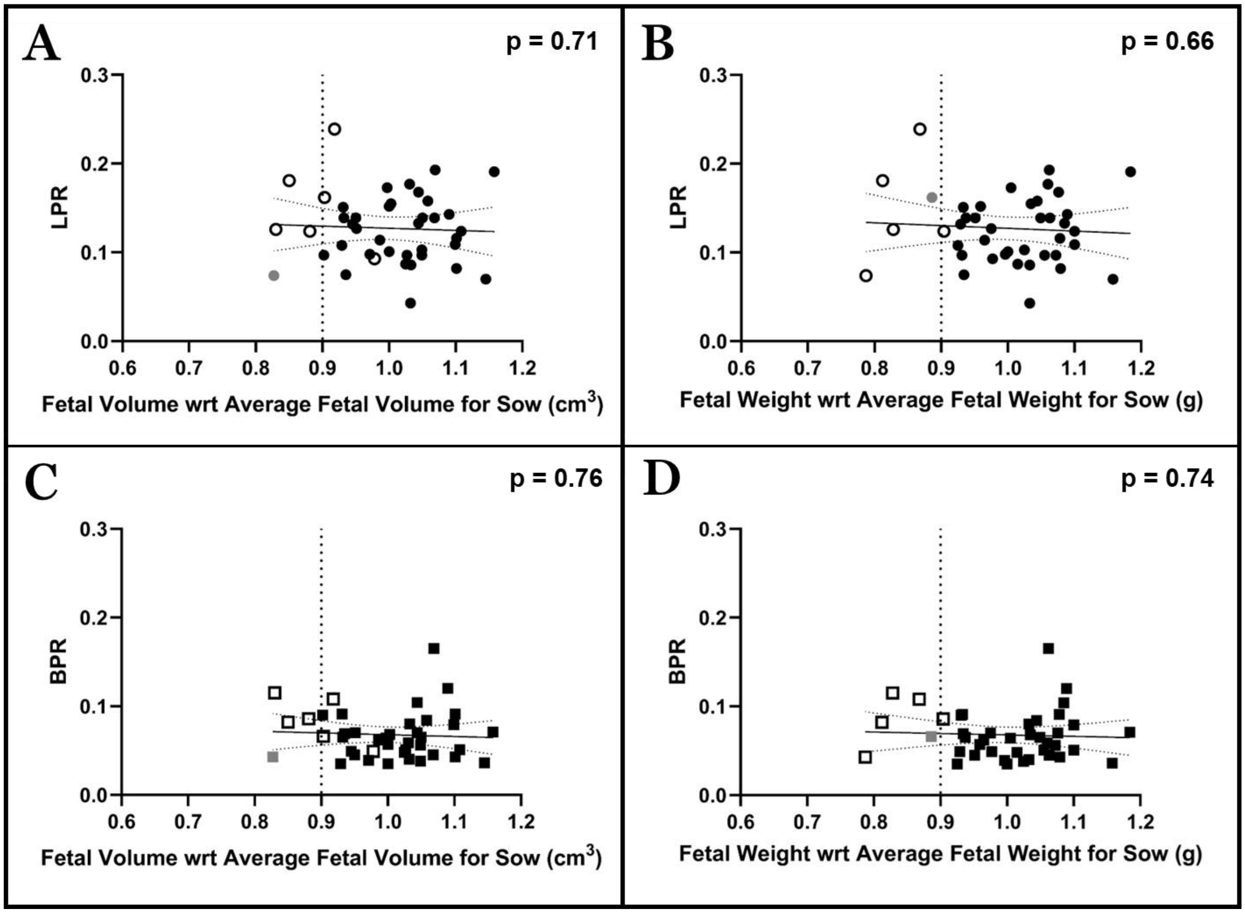


**Figure S1: Linear regression for (A) lactate pyruvate ratio (LPR: circles) and fetal volume ($cm^3$) / average fetal volume in pregnancy, (B) LPR and fetal weight (g) / average fetal weight in pregnancy, (C) bicarbonate pyruvate ratio (BPR: squares) and fetal volume ($cm^3$) / average fetal volume in pregnancy, (D) BPR and fetal weight (g) / average fetal weight in pregnancy, with 95% confidence intervals. The dotted line for each represents the cut-off value for the spontaneous intrauterine growth restricted (spIUGR) marker (0.9 $cm^3$ or g). Full black colour: non-spIUGR fetuses, outline black colour: spIUGR fetuses (three or more markers for spIUGR), and full gray: non-spIUGR fetuses that had that marker as spIUGR. The metabolite alanine (alanine pyruvate ratio (APR)) was not included due to low signal-to-noise in the placenta.**

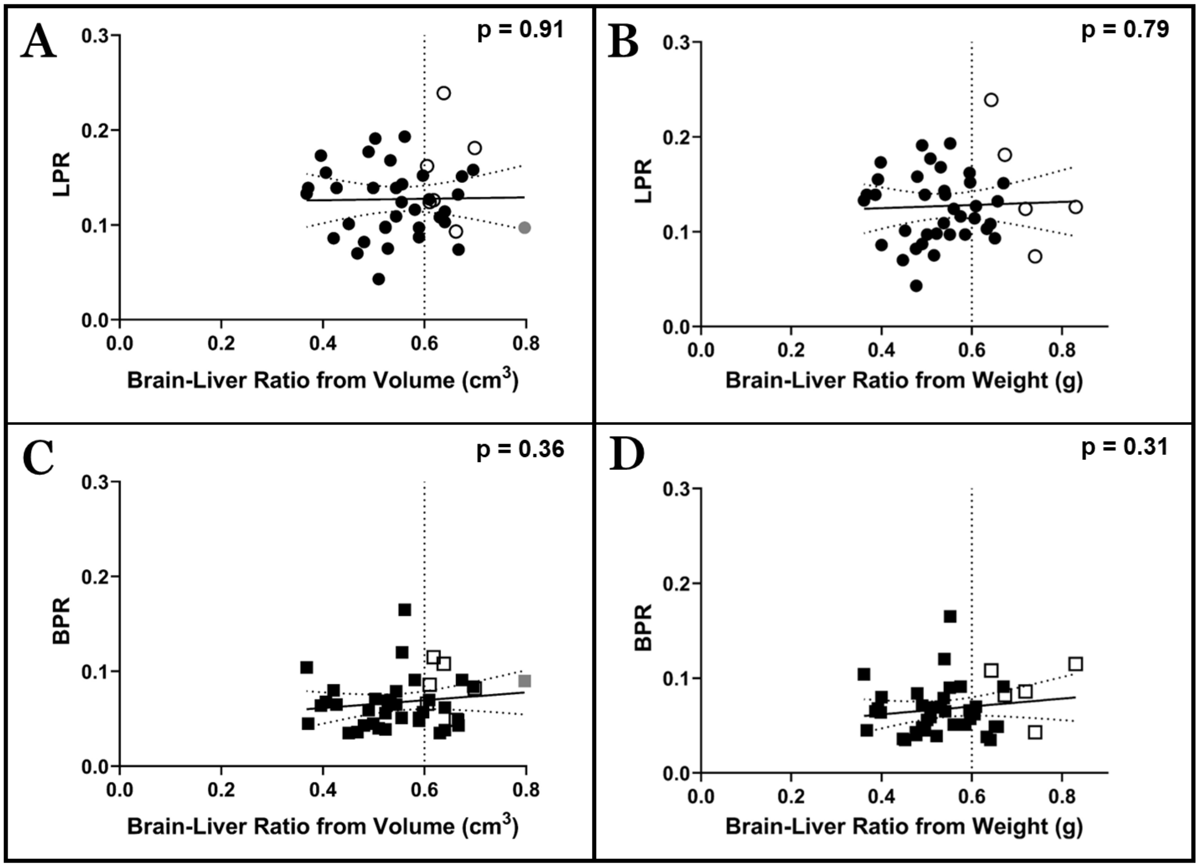


**Figure S2: Linear regression for (A) lactate pyruvate ratio (LPR: circles) and brain-liver volume ($cm^3$) ratio, (B) LPR and brain-liver weight (g) ratio, (C) bicarbonate pyruvate ratio (BPR: squares) and brain-liver volume ratio, and (D) BPR and brain-liver weight ratio, with 95% confidence intervals. The dotted line for each represents the cut-off value for the spontaneous intrauterine growth restricted (spIUGR) marker (0.06 $cm^3$ or g). Full black colour: non-spIUGR fetuses, outline black colour: spIUGR fetuses (three or more markers for spIUGR), and full gray: non-spIUGR fetuses that had that marker as spIUGR. The metabolite alanine (alanine pyruvate ratio (APR)) was not included due to low signal-to-noise in the placenta.**

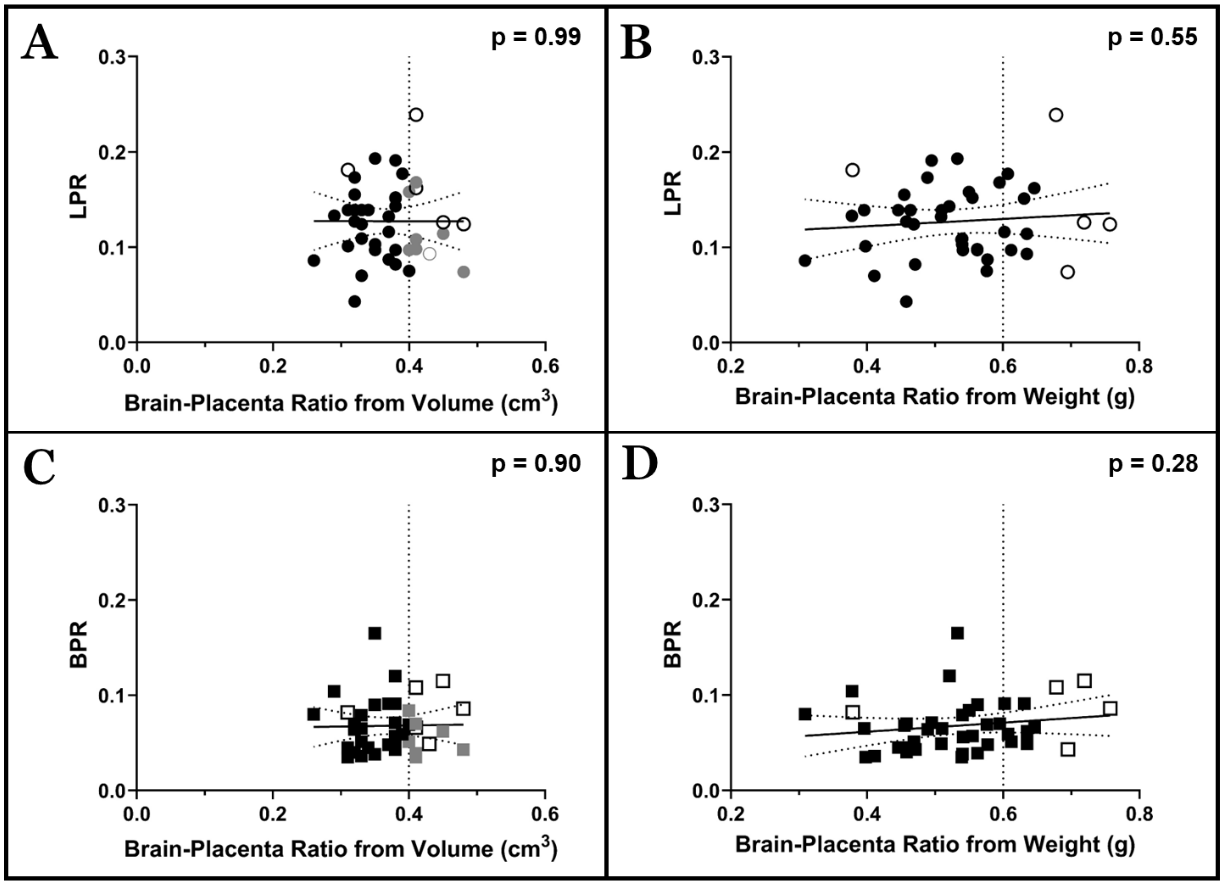


**Figure S3: Linear regression for (A) lactate pyruvate ratio (LPR: circles) and brain-placenta volume ($cm^3$) ratio, (B) LPR and brain-placenta weight (g) ratio, (C) bicarbonate pyruvate ratio (BPR: squares) and brain-placenta volume ratio, and (D) BPR and brain-placenta weight ratio, with 95% confidence intervals. The dotted line for each represents the cut-off value for the spontaneous intrauterine growth restricted (spIUGR) marker (0.04 $cm^3$ or 0.06 g). Full black colour: non-spIUGR fetuses, outline black colour: spIUGR fetuses (three or more markers for spIUGR), and full gray: non-spIUGR fetuses that had that marker as spIUGR. The metabolite alanine (alanine pyruvate ratio (APR)) was not included due to low signal-to-noise in the placenta.**

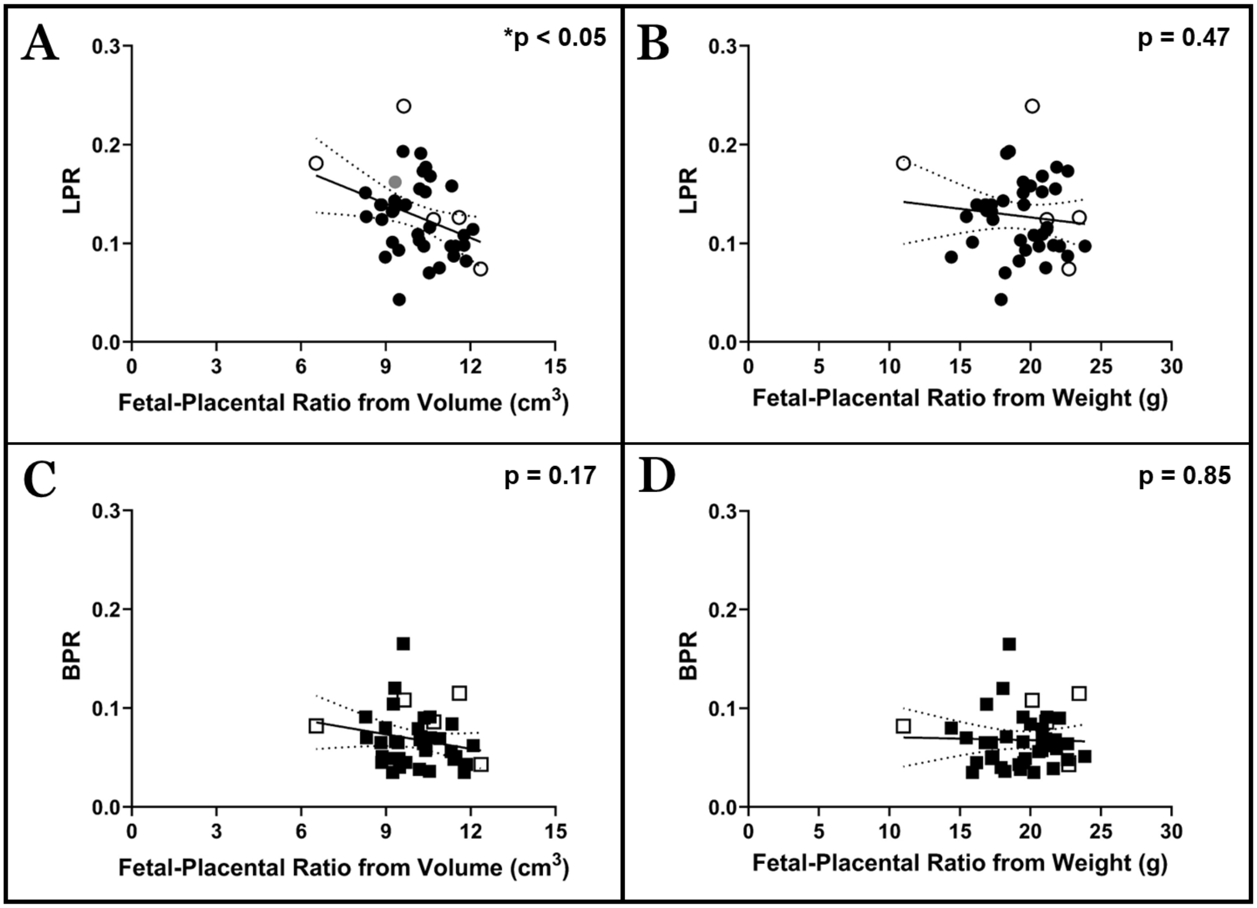


**Figure S4: Linear regression for (A) lactate pyruvate ratio (LPR: circles) and fetal-placental volume ($cm^3$) ratio, (B) LPR and fetal-placenta weight (g) ratio, (C) bicarbonate pyruvate ratio (BPR: squares) and fetal-placenta volume ratio, and (D) BPR and fetal-placenta weight ratio, with 95% confidence intervals. Full black colour: non-spIUGR fetuses, outline black colour: spIUGR fetuses (three or more markers for spIUGR), and full gray: non-spIUGR fetuses that had that marker as spIUGR. The metabolite alanine (alanine pyruvate ratio (APR)) was not included due to low signal-to-noise in the placenta.**
